# Hippocampus-sensitive and striatum-sensitive learning one month after morphine or cocaine exposure in male rats

**DOI:** 10.1101/2021.11.07.467628

**Authors:** Robert S. Gardner, Donna L. Korol, Paul E. Gold

**Affiliations:** Department of Biology, Syracuse University, Syracuse, NY, 13244; Now at Upstate Medical University, Department of Neurosurgery, Syracuse NY 13210

**Author notes:** Corresponding Author: Paul E Gold, Syracuse University, Department of Biology, Life Science Complex, Syracuse, NY, 13244.

**Keywords:** Decision-making, Learning, GFAP, Astrocytes, BDNF, GSK3β

## Abstract

These experiments examined whether exposure to drugs of abuse altered the balance between hippocampal and striatal memory systems as measured long after drug treatments. Male rats received injections of morphine (5 mg/kg), cocaine (20 mg/kg), or saline for five consecutive days. One month later, rats were then trained to find food on a hippocampus-sensitive place task or a striatum-sensitive response task. Relative to saline controls, morphine-treated rats exhibited impaired place learning but enhanced response learning; prior cocaine exposure did not significantly alter learning on either task. Another set of rats was trained on a dual-solution T-maze that can be solved with either place or response strategies. While a majority (67%) of control rats used place solutions in this task, morphine treatment one month prior resulted in a shift to response solutions exclusively (100%). Prior cocaine treatment did not significantly alter strategy selection. Molecular markers related to learning and drug abuse were measured in the hippocampus and striatum one month after drug exposure in behaviorally untested rats. Protein levels of glial-fibrillary acidic protein (GFAP), an intermediate filament specific to astrocytes, increased significantly in the hippocampus after morphine, but not after cocaine exposure. Exposure to morphine or cocaine did not significantly change levels of brain-derived neurotrophic factor (BDNF) or a downstream target of BDNF signaling, glycogen synthase kinase 3β (GSK3β), in the hippocampus or striatum. Thus, exposure to morphine results in a long-lasting shift from hippocampal toward striatal dominance during learning. The effects of prior morphine injections on GFAP suggest that long-lasting alterations in hippocampal astrocytes may be associated with these behavioral strategy shifts.

## 1. Introduction

This report applies a multiple memory systems perspective to evaluate persistent behavioral and brain changes after exposure to cocaine or morphine. Many regions of the brain differentially contribute to learning and memory in tasks with distinct attributes (McDonald and White, 1994; Hunsaker and K.esner, 2018). Of note here, the hippocampus contributes importantly to spatial or contextual learning and memory, and goal-directed decision-making while the dorsolateral striatum is important for learning based on stimulus-response associations and the formation and expression of well-learned behaviors or habits (Gold et al., 2013; Goodman and Packard, 2016; Korol and Pisani, 2015; Poldrack and Packard, 2003; White et al., 2013).

One characteristic of multiple memory systems is that they sometimes compete for control over the expression of learning. For example, impairments of hippocampal functions can enhance striatum-sensitive learning while impairments of striatal functions can enhance hippocampus-sensitive learning (e.g., Chang and Gold, 2003; Gardner et al., 2020a; Lee et al., 2008; McElroy and Korol, 2005; Mitchell & Hall, 2008). Moreover, drugs of abuse may influence the balance of functions across multiple memory systems with effects that could lead to increased drug seeking (Goodman and Packard, 2016; Lovinger and Gremel, 2020; White, 1996). Notably, acute exposure to alcohol facilitates the use of dorsolateral striatum cue or response strategies over spatial or place strategies (Matthews et al., 1999; Sun et al., 2018). If drugs of abuse create an imbalance across systems to favor reliance on a dorsolateral striatal system, drug exposure may likewise enhance the use of habits to guide behavior. A chronic change in competitive interactions across memory systems could contribute to persistent use of striatal-based habit strategies, with the transition from occasional to habitual drug-seeking behaviors a consequence of this functional imbalance across systems.

Most studies of drug-induced changes in learning and memory have been conducted in rats or mice while under the influence of the drug or soon after withdrawal. However, relapse following long-term abstinence is common in addiction, an observation consistent with the hypothesis that functional imbalances across learning and decision-making systems persist long after drug use. Importantly, studies that examined learning and memory after prolonged periods of abstinence have primarily assayed a single memory system, most often the hippocampal system, and have not assessed possible drug-induced shifts across multiple systems. For example, in rodents abstinent for 1-3 months, cocaine impaired learning and memory in the spatial version of the swim task (Mendez et al., 2008), object location recognition, and spatial working memory (Guevara-Miranda et al., 2017). Morphine exposure impaired performance on a spatial alternation task after four weeks of abstinence (Shahroodi et al, 2020) and learning on a food-rewarded conditioned place preference task after five weeks of abstinence (Harris and Aston-Jones, 2003). Interference with hippocampal functions impairs learning on each of these paradigms supporting their designation of hippocampus-dependent tasks (Ferbinteanu and McDonald, 2001; Gerlai, 2001; Hitchcock and Lattal, 2018; Kesner, 2018; Korol et al., 2019).

The primary focus of this report was to determine whether a 5-day treatment of cocaine or morphine differentially altered hippocampal and striatal memory system functions after one month of abstinence. The drugs were selected on the basis of important societal abuse of each drug coupled with their disparate cellular mechanisms of action. Separate sets of rats were trained on single-solution place and response mazes for which hippocampal or striatal strategies, respectively, provide effective solutions. A third set of rats was trained on a dual solution T-maze in which either place or response strategies afford effective solutions; probe tests administered after dual-solution training identify the dominant strategy used on the task (Restle, 1957; Tolman et al., 1946). These tasks have been used reliably to reveal changes across multiple memory system functions resulting from extensive training, advanced age, stress, and hormone status (e.g., Gardner et al., 2013, 2016, 2020a,b; Gold et al., 2013; Gold, 2016; Hawley et al., 2012; Korol and Pisani, 2015; Korol and Wang, 2018; Packard and Goodman, 2012, 2013; Packard et al., 2018; Packard and McGaugh, 1996; Schwabe, 2013; Zurkovsky et al., 2007).

Additional experiments measured tissue levels of glial fibrillary acidic protein (GFAP), brain-derived neurotrophic factor (BDNF), and glycogen synthase kinase-3 beta (GSK3β) across brain regions one month after cocaine or morphine exposure. GFAP was selected as a marker of astrocytic responses to treatments and was tested here based on evidence that astrocytes are important in the regulation of learning and memory (Alberini et al., 2018; Gao et al., 2016; Gold, 2014; Hertz and Chen, 2018; Magistretti, 2006; Newman et al., 2011; Pellerin and Magistretti, 2012; Santello et al., 2019; Steinman et al., 2016; Suzuki et al., 2011), including findings that astrocytic actions dissociate by memory system (Gold, 2016; Korol et al., 2019; Newman et al., 2017; Scavuzzo et al., 2021). Astrocyte functions are also important in the contexts of drug abuse and relapse (e.g., Boury-Jamot et al., 2016; Haydon et al., 2009; Lindberg et al., 2019; Ozawa et al., 2001; Song and Zhao, 2001; Zhang et al., 2016). BDNF, through activation of its receptor target tropomyosin receptor kinase B (TrkB), supports learning, memory, and synaptic plasticity in hippocampal and striatal systems (e.g., Andero et al., 2014; Bekinschtein et al., 2014; Cunha et al., 2010; Korol et al., 2013; Lu et al., 2008; Miranda et al., 2019). Pharmacological manipulations of BDNF and the TrkB receptor modulate drug-seeking behaviors and BDNF levels in different brain regions change after drug experience (e.g., Geoffroy and Noble, 2017; Harvey et al., 2019; Li & Wolf, 2015; Shahroodi et al., 2020), pointing to the role of BDNF signaling in learning and drug exposure. Downstream of TrkB activation, a PI3K-Akt pathway modulates GSK3β phosphorylation at serine 9; GSK3β is a molecular target with a role both in learning and memory and drug abuse (e.g., Dewachter et al 2009; Eagle and Robison, 2018; Miller et al., 2014). Thus, the molecular markers selected are each associated both with learning and memory and with drugs of abuse and may therefore reflect drug-induced changes in learning abilities across memory systems.

## 2. Methods

### 2.1 Animals

Male Sprague Dawley rats (N = 117) were obtained from Envigo (Indianapolis, IN). The rats were three months old at the time of arrival at Syracuse University and were singly housed in a 12:12 hr light/dark cycle (7:00 a.m. lights on, 7:00 p.m. lights off) with free access to food and water until initiation of food restriction ∼10 days before maze testing. All methods were carried out in accordance with the National Institutes of Health Guide for the Care and Use of Laboratory Animals and were approved by the Syracuse University Institutional Animal Care and Use Committee.

### 2.2 Drug injections and food restriction

The overall experimental design is presented in Fig. 1. On each of five consecutive days, rats received an injection (i.p., 1 ml / kg) of either 0.9% saline, cocaine (20 mg / kg; Sigma-Aldrich, St. Louis, MO), or morphine (5 mg / kg; Sigma). Drug doses were chosen to match those shown to support conditioned place preference (Denny and Unterwald, 2019; White et al., 2005). Cocaine and morphine exposure induced modest weight loss during the five-day injection period while the saline rats increased their weight at that time (weight changes: saline, +4.3 ± 0.8 gm; cocaine: -3.3 ± 1.5 gm; morphine: -1.9 ± 1.1 gm). Body weights returned to control weights thereafter and were equivalent across groups at the start of food restriction (F_2,109_ = 0.9; p > 0.40; data not shown). Twenty days after the final (fifth) injection, rats were placed on food restriction for ∼10 days to reduce body weights to 80-85% of their free-feeding weights. During food restriction, rats were also handled each day for several minutes and given several small pieces of Frosted Cheerios^®^ cereal, the reward used during training. The effects of drug exposure on levels of select protein targets were assessed in separate groups of untrained food-restricted rats.

**Fig. 1.**
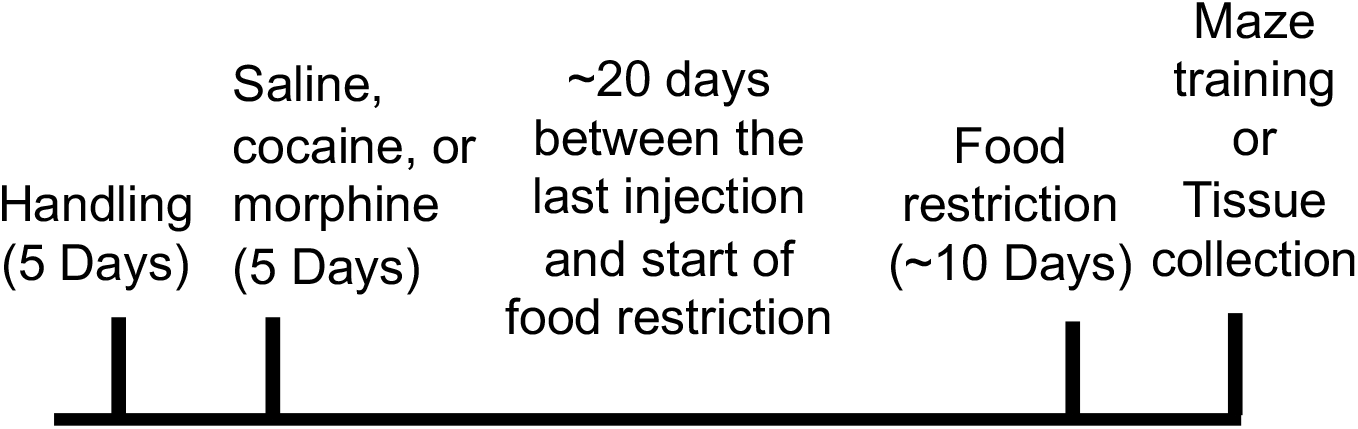
Experimental design. Rats were handled for five days prior to receiving once daily i.p. injections (1 mg / kg) of 0.9% saline, cocaine (20 mg / kg), or morphine (5 / mg / kg) for five days. After food restriction, rats were trained on one of three maze tasks one month after the last injection. A separate group of untrained rats was euthanized one month after the last drug injection and hippocampal and striatal tissue were collected and subsequently assayed for levels of glial fibrillary acidic protein, brain-derived neurotrophic factor, and GSK3β.

### 2.3 Training

Approximately one month after drug treatments (mean ± SD = 33 ± 2 days), rats were trained in a single session on either a place, response, or dual-solution task (Fig. 2 A-C) with training beginning between 4-8 hrs into the light cycle; each rat was trained on only one of the three tasks. Before training, each rat was placed in a clean cage and brought into the training room to acclimate for 15 minutes. Rats were trained to find food (∼1/2 Frosted Cheerios^®^ piece) on a plus-shaped maze configured into a T-maze by blocking one arm. The maze arms measured 45 × 14 × 7.5 cm (length x width x wall height) with a 14 × 14 cm center area. The goal cups at the ends of each possible goal arm contained inaccessible pieces of reward to minimize the use of navigation based on olfaction.

**Fig. 2.**
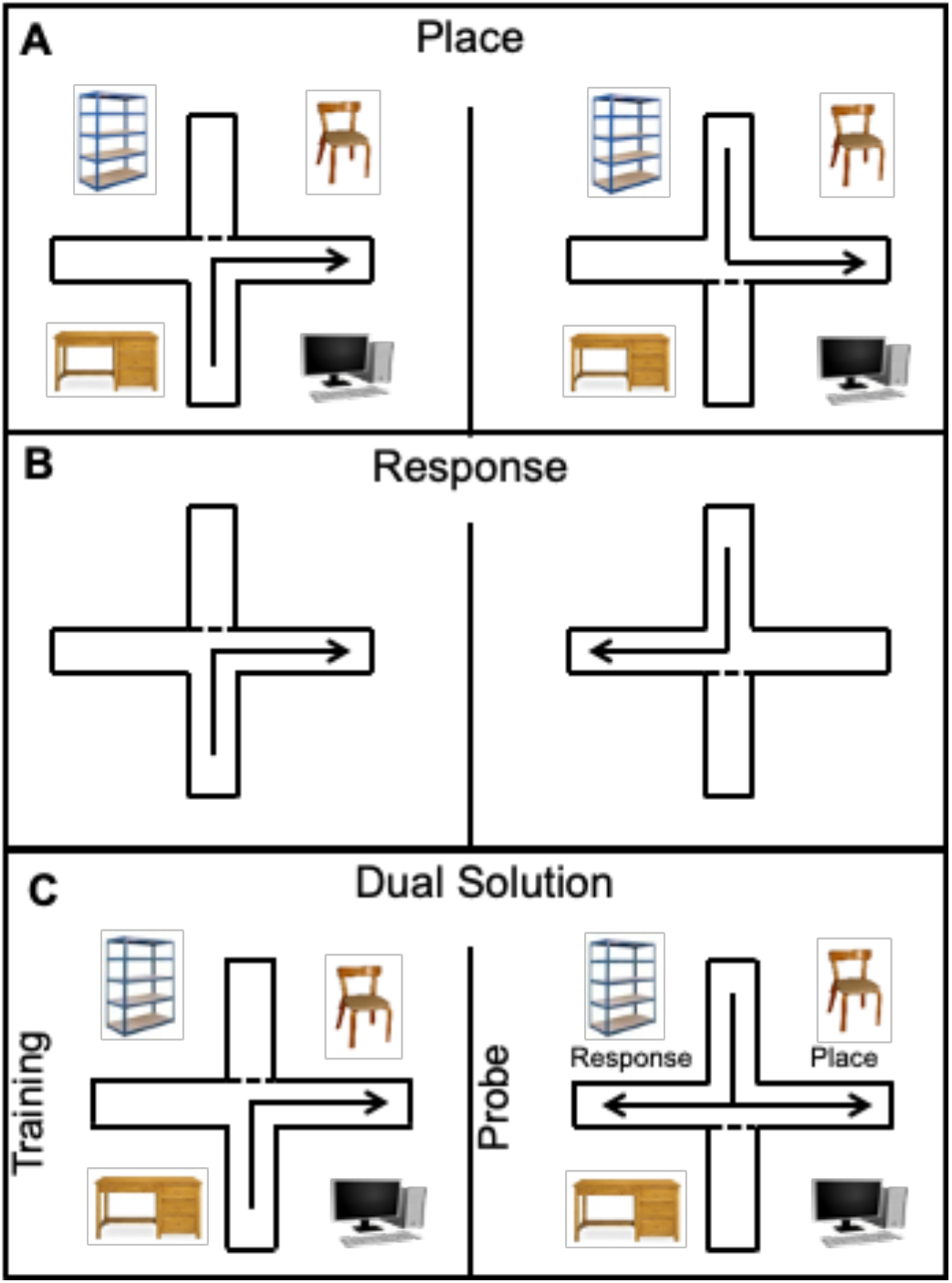
Training tasks. (A) Place learning involved finding the food reward in a specific room position. (B) Response learning used a fixed body turn to find the food reward. (E) A dual-solution task could be solved using either place or response strategies; after learning (9/10 correct trials), the rat was placed in the opposite start position to that used during training on each of three probe trials. The arm choice made on two of three or three of three probe trials identified the dominant learning strategy: place or response.

Training on the place task (100 trials, 30 s inter-trial interval) was conducted in a room with a rich array of extra-maze cues that could be used to locate the position of the goal arm (Fig. 2A). On each trial, a rat was started from one of two possible start positions. The Frosted Cheerios^®^ reward was placed in the goal arm located in the same room position across trials. Training on the response task (75 trials, 30 s inter-trial interval) was conducted in a room with minimal extra-maze cues; a solid beige curtain surrounded the maze. On each trial, a rat was started from one of two possible start positions. The rat was rewarded for making a consistent turn (e.g., turn right) from either start arm (Fig. 2B). For both tasks, start arms were semi-randomly assigned with no more than 3 consecutive starts from the same arm. Goal arm (east vs. west for Place) and rewarded turn (right vs. left for Response) were randomly assigned and counterbalanced across all rats within treatment condition. The experimental design included three groups (saline, morphine, cocaine) of 10 rats each for place and response tasks. However, during place training, one rat had a strong turning bias making the same turn on 46 of the first 50 trials and thus its data were excluded from analyses.

Training on the dual-solution task was conducted in a room with extra-maze cues (Fig. 2C). The start arm and position of the reward were kept constant across the training session such that use of either a place or a response strategy resulted in a “correct” rewarded choice. The rewarded arm was counterbalanced across all rats within treatment condition. After a rat reached 9/10 correct trials, the rat was given a series of three probe trials to identify the dominant strategy controlling behavior. On probe trials, the rat was started in the arm opposite to that used during training. If the rat entered the arm in the same room position as rewarded during the training trials, the rat was identified as using a place strategy. If the rat made the same turn as during training, it was identified as using a response strategy. As neither choice was “incorrect,” both arms were baited on probe trials. A single training trial was administered from the original training start arm between each probe trial. Rats were grouped into those using place or response strategies based on the strategy expressed on the majority (2/3) of their probe trials, as done previously (Gardner et al., 2020b). On the dual-solution task, two rats in the cocaine group and one rat in the morphine group reached the 9/10 learning criterion within the first ten trials, thereby exhibiting a directional bias that did not likely reflect learning. One rat in the morphine group failed to reach the learning criterion during the dual-solution training session, precluding strategy assessment. The data from these rats were excluded from further analyses. Final sample sizes for the dual-solution task were 9, 8, and 9 for saline, cocaine, and morphine conditions, respectively.

### 2.4 Tissue collection

Protein levels of GFAP, BDNF, and GSK3β in hippocampus and striatum were assessed in untrained rats one month after drug exposure (n = 27). To match the design of the behavioral studies above, these rats underwent food-restriction during the 10 days prior to sample collection. Rats received an overdose of sodium pentobarbital (50 mg/rat) one month after drug exposure. Their brains were excised and the hippocampi and dorsal striata were dissected and frozen on dry ice. Frozen samples were pulverized on dry ice using a mortar and pestle, aliquoted for subsequent assays, and stored at -80°C.

### 2.5 ELISAs for quantification of GFAP and BDNF expression

Hippocampal and striatal samples were suspended in homogenization buffer (∼14 mg tissue / ml buffer; 100 mM PIPES buffer [pH 7.0], 500 mM NaCl, 2 mM EDTA, 200 µM PMSF, 1 µM pepstatin, 10 µM leupeptin, 0.3 µM aprotenin, 0.2% Triton-X-100) and homogenized (Polytron PT 1600 E, Kinematica; max speed) for ∼10 s on ice. Samples were centrifuged at 16,000g for 30 min at 4°C. Pro and mature forms of BDNF were measured in the supernatant with sandwich ELISA kits (Biosensis BEK-2217 and BEK-2211; Thebarton, South Australia) using 4x and 5x dilutions, respectively. GFAP was also measured in the supernatant with a sandwich ELISA kits (Sigma NS830) using a 200x dilution. Concentrations of target proteins were computed using standard curves and normalized to the respective weight of the tissue sample measured prior to homogenization.

### 2.6 Semiquantitative western blotting for GSK3β and pGSK3β

Hippocampal and striatal samples were suspended in homogenization buffer (∼20 mg tissue / 80µl buffer; 1 mM EGTA, 1 mM EDTA, 20 mM Tris [pH 7.4], 1 mM sodium pyrophosphate tetrabasic decahydrate, protease cocktail inhibitor [Roche, Mannheim, Germany], phosphatase cocktail inhibitor [Phosstop; Roche]) and homogenized (Polytron PT 1600 E; max speed) for ∼10 seconds on ice. Samples were centrifuged at 5700g for 5 min at 4°C. Total protein was assayed in the supernatant using the Pierce BCA assay. The supernatant was subsequently diluted and mixed with Protein Loading Buffer (LI-COR Biosciences; Lincoln, NE) containing 10% beta-mercaptoethanol and heated to 95°C degrees for 10 min. A total protein concentration of 3 µg / µl was used for western blot experiments.

Experimental samples (20 µg total protein, 6.67 µl) were loaded onto 10% SDS-polyacrylamide gels and resolved via electrophoresis for 60 min at 200 V. Gels were run separately for each brain region. Samples were balanced with respect to drug condition within each gel. Pooled hippocampal or striatal homogenates were also loaded at three total protein amounts (10, 20, and 30 µg) to test linearity of immunoblotting and to normalize signals across runs. All Blue Prestained Protein Standards (1-3 µl; BioRad Laboratories, Hercules, CA) were used to identify band molecular weights. Following electrophoresis, proteins were transferred (100 V, 2 hr, 4°C) to polyvinylidene difluoride membranes (Immobilon-FL, Millipore, Billerica, MA). To confirm protein transfer, membranes were stained in Ponceau S Solution (1 min; Sigma) and washed in H_2_O followed by tris-buffered saline (TBS; 20 mM Tris, 150mM NaCl, pH 7.5). Membranes were blocked for 1 hr in TBS Odyssey blocking buffer (LI-COR) and incubated at 4°C for 18-22 hr in a cocktail comprising anti-GSK3β mouse monoclonal antibody (1:1500; #9832, Cell Signaling, Rockford, IL), anti-phospho GSK3β (Ser9) rabbit monoclonal antibody (1:1000; #9323, Cell Signaling), and anti-β-tubulin mouse monoclonal antibody for loading control (1:30,000; #86298T, Cell Signaling). Subsequently, membranes were washed 4 times (10 min each) in TBS-Tween (0.1% Tween 20) and incubated (1 hr) with IRDye 800 CW goat anti-rabbit (1:10000; LI-COR Biosciences) and IRDye 680 RD goat anti-mouse (1:10000; LI-COR 129 Biosciences) antibodies in blocking buffer (0.1% Tween 20; 0.1% SDS). Membranes were again washed 4 times in TBS-Tween, followed by a rinse (2 min) in H_2_O.

Blots were scanned on an Odyssey near-infrared imaging system (LI-COR Biosciences) and analyzed for relative levels of GSK3β, pGSK3β, and β-tubulin using Image Studio software (LI-COR Biosciences). After target bands were identified and fitted manually, integrated optical density (OD) was automatically corrected for background (median OD 3 pixels above and below each band) using the Studio software. Several steps were taken to quantify relative protein levels reliably as follows. It was ensured that all blots displayed linear curves from pooled standard samples. To account for loading variability, GSK3β and pGSK3β ODs were normalized to ODs from β-tubulin. To account for between-blot variability, an additional normalization to the back-calculated 20-µg pooled standard sample was made. Finally, averages of each sample, run in duplicate on different blots, were used for summary data.

### 2.7 Data analysis and statistics

For place and response tasks, learning was quantified as both the number of trials needed to reach a learning criterion and as trial accuracy across training. Trials to criterion (TTC) was set at nine correct trials of the most recent 10, with at least 6 consecutive trials correct in that span, and trial accuracy was measured as percent correct across training. Two-way ANOVAs assessed interaction effects of drug treatment and task on TTC and trial accuracy. ANOVAs run separately for place and response tasks tested differences in these measures across drug groups. The Fisher Least Significant Differences (LSD) method was used to make planned pairwise comparisons.

For the dual-solution results, chi square tests were used to determine differences in strategy use among cocaine and morphine rats compared to strategy use in saline control rats. An ANOVA with the LSD method applied to planned comparisons was used to assess differences in TTC across drug conditions in the dual-solution task.

One-way ANOVAs with the LSD method for planned comparisons were used to assess differences across drug conditions in body weights. A two-way ANOVA tested interaction effects between drug treatment and brain region on levels of GFAP, proBDNF, mature BDNF (mBDNF), total GSK3β, pGSK3β, and the ratio of phosphorylated to total GSK3β; one-way ANOVAs with the LSD method were used for planned comparisons within each brain region. Assays for BDNF used a subset of collected brain samples (n = 22). Three protein samples were lost because of technical issues, one from each drug condition. The final Ns in each assay are shown in the respective figure legends.

Statistical analyses were performed using Prism 9 for macOS and SPSS, with α = 0.05.

## 3. Results

Morphine treatment had opposite effects on place and response learning, impairing place learning but enhancing response learning. Cocaine treatment, however, did not significantly affect learning in either task (Fig. 3A-F).

**Fig. 3.**
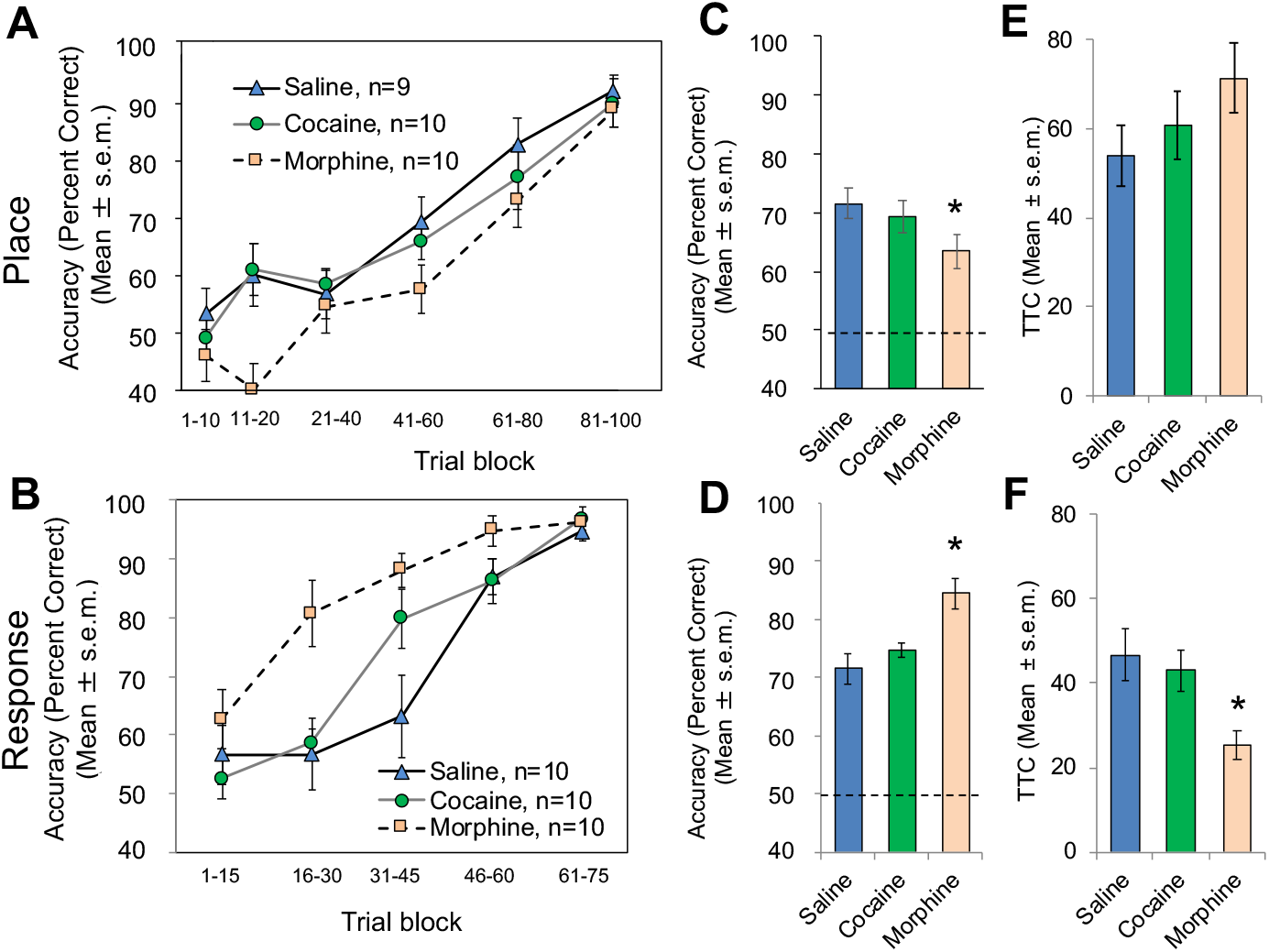
Cocaine and morphine effects on place and response learning tested one month after drug exposure. Changes in trial accuracy throughout the training sessions are shown for (A) place and (B) response tasks. Accuracy collapsed across all trials (100 for place and 75 for response) is shown for (C) place and (D) response tasks. Dashed lines indicate chance levels (50%). The number of trials to criterion (TTC; 9/10 correct trials with at least 6 consecutively correct) is shown for (E) place and (F) response tasks. Note that prior morphine exposure resulted in reduced accuracy and a trend toward increased trials to criterion on the place task. In marked contrast, morphine resulted in increased accuracy and decreased trials to criterion on the response task. Prior cocaine exposure did not significantly affect place or response learning measures. * p < 0.05 vs. saline. Place: Saline n = 9, Cocaine n = 10, Morphine n = 10; Response: Saline n = 10, Cocaine n = 10, Morphine n = 10.

**Fig. 4.**
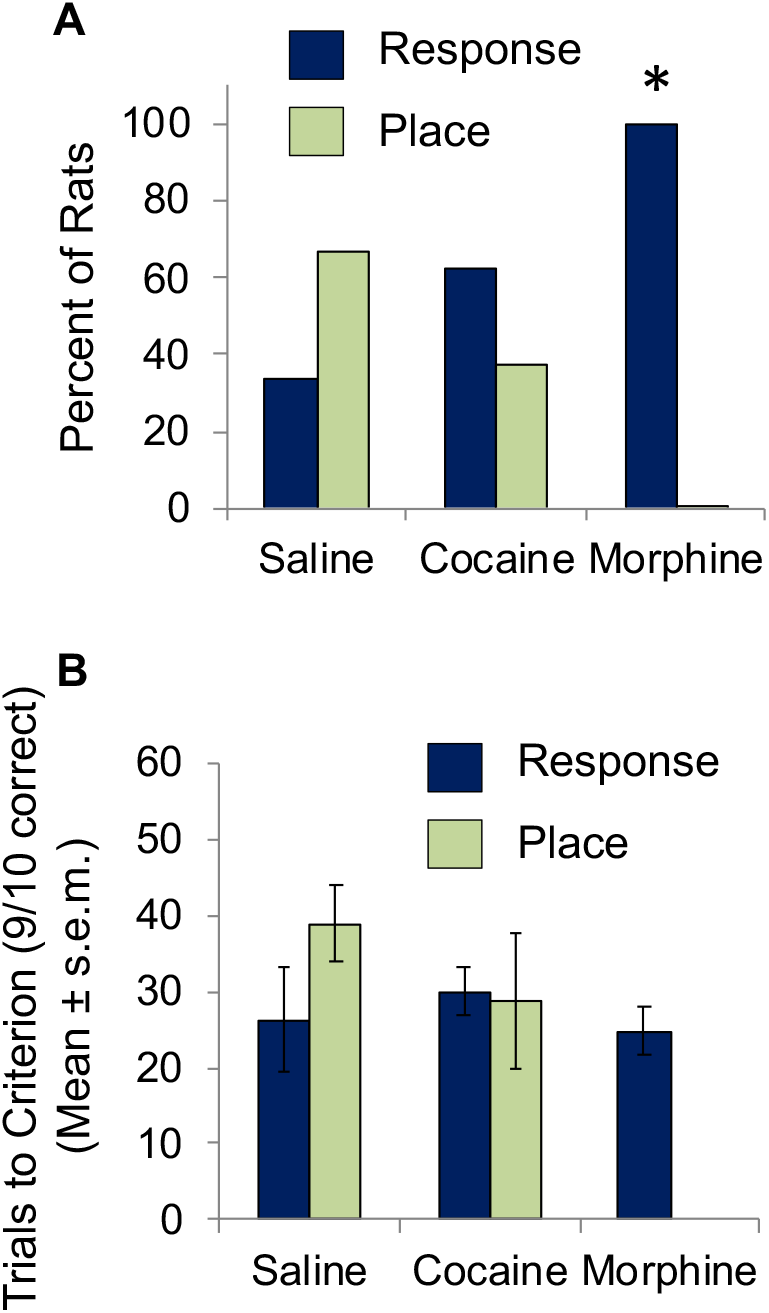
Cocaine and morphine effects on dual-solution strategies tested one month after drug exposure. (A) The percent of rats expressing place vs. response strategies on probe trials shifted from 67% place in the saline group to 100% response after prior exposure to morphine. The strategies expressed by rats after cocaine showed a modest shift in expression of place to response strategies that did not differ significantly from the strategies used by the saline group. (B) The number of trials to reach the 9/10 learning criterion (TTC) was comparable across conditions. * p < 0.05 morphine vs. saline. Saline n = 9, Cocaine n = 8, Morphine n = 9.

Morphine significantly impaired place learning as measured by accuracy across all trials (p < 0.05; Fig. 3C), particularly during the early trials. A trend for morphine-induced impairment was also observed on the TTC measure, although this difference did not reach statistical significance (p > 0.10; Fig. 3E). Conversely, learning on the response task was significantly enhanced on both accuracy and TTC measures in morphine-treated rats compared to controls (p’s < 0.05; Fig. 3D, 3F). The overall enhancement of response learning by morphine reflects an increased accuracy early in training in morphine vs. saline groups (Fig. 3B). Differential effects of morphine on the two tasks were confirmed by a significant task by drug interaction on both trial accuracy (F_1,35_ = 15.63, p < 0.001) and TTC (F_1,35_ = 9.57, p < 0.01); the main effect of drug for accuracy (F_1,35_ = 0.84, p > 0.3) or TTC (F_1,35_ = 0.09, p > 0.7) was not significant, reflecting the opposite direction of drug effects on place and response learning.

In contrast, cocaine treatment did not produce significant effects on either place or response learning (Fig. 3C-3F). Planned comparisons between cocaine vs. saline groups using both accuracy and TTC measures of learning on each task revealed p’s > 0.35 for all tests. The absence of an effect of cocaine on later learning was also apparent by the absence of a significant interaction of task by drug on either accuracy (F_1,35_ = 1.32, p > 0.2) or TTC (F_1,35_ = 0.68, p > 0.4).

Of note, choice accuracy was near chance (50%) at the onset of training across drug conditions in both the place and response tasks. Accuracy on trials 1-10 for place learning and trials 1-15 for place learning did not differ by drug condition (all p’s > 0.1; Figs. 3A, 3B).

In the dual-solution task, we observed a robust shift away from the use of place strategies toward reliance on response strategies after exposure to morphine. All rats that had received morphine one month prior to training expressed response solutions in this task as compared to only a third of saline controls, a difference that was statistically significant (Fig. 3A; X^2^ = 9.0, p < 0.001). Cocaine-treated rats showed a modest shift toward the use of response strategies, with 62.5% expressing response solutions. However, differences in strategy use by saline-treated vs. cocaine-treated rats was not significant (X^2^ = 1.45, p > 0.20). The number of trials rats took to reach the 9/10 learning criterion on the dual-solution task was consistent across drug conditions regardless of strategy used during the probe trials (F_2,23_ = 1.89 p > 0.10; Fig. 3B).

GFAP expression was increased in the hippocampus of untrained rats one month after morphine exposure when compared to that seen in saline controls (morphine vs. saline: p < 0.05; main effect of drug: F_2,23_ = 2.8, p < 0.09; Fig. 5). Hippocampal GFAP in cocaine-treated rats, although elevated, was not significantly above that of saline-treated rats (p >0.10). Striatal GFAP was comparable across drug conditions (F_2,24_ = 0.3, p > 0.7; morphine or cocaine vs. saline: p’s > 0.40). No interaction was found between brain region and drug condition on levels of GFAP (F_2,47_ = 2.0, p > 0.1). There was a significant main effect of brain region on GFAP content (F_1,47_ = 177.7, p < 0.0001) with higher levels in the hippocampus than in the striatum.

**Fig. 5.**
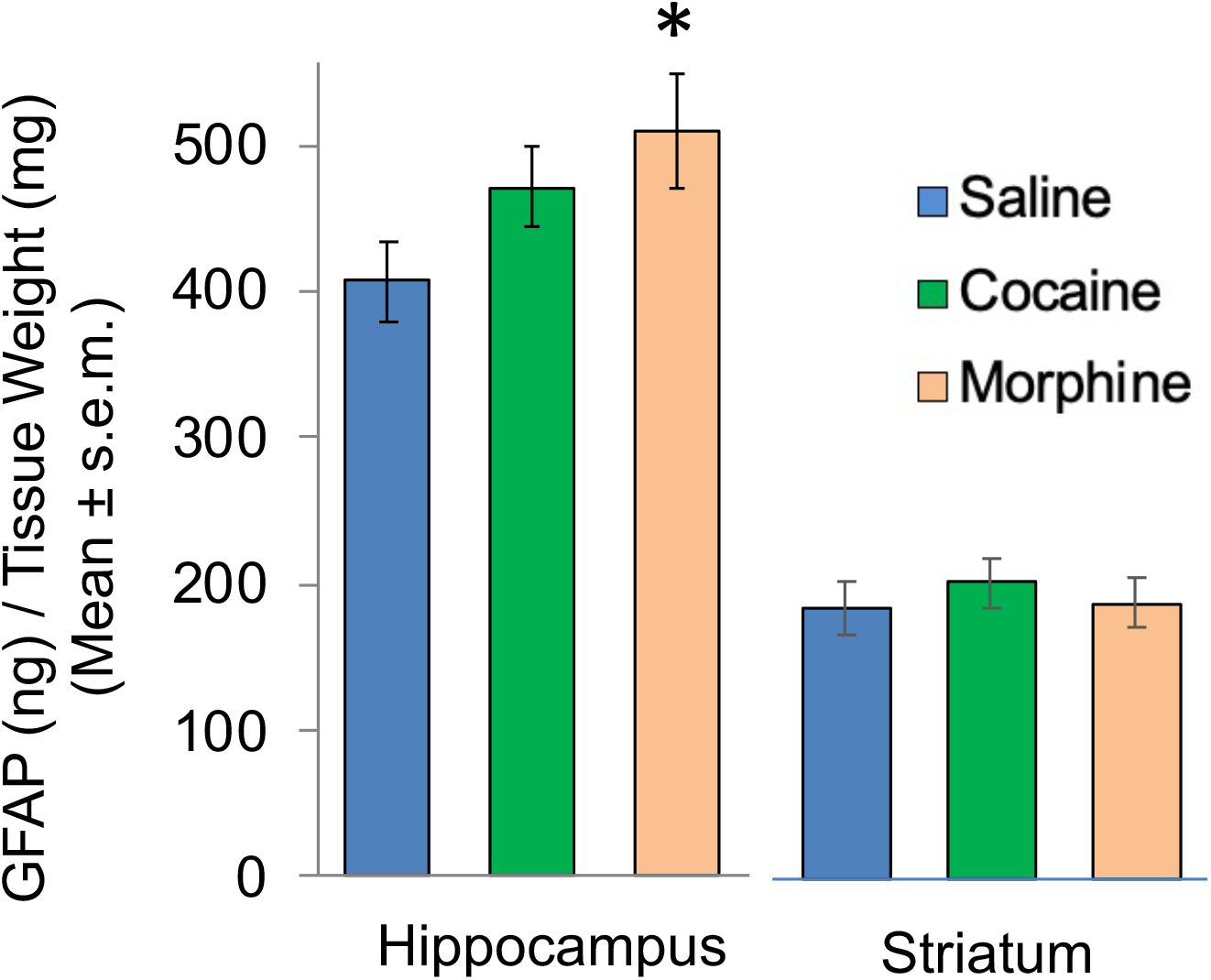
Glial fibrillary acidic protein (GFAP) levels (ng/mg wet weight) in the hippocampus and striatum one month after drug exposure as measured with ELISAs. Note that GFAP levels increased in the hippocampus but not in the striatum after morphine treatment. GFAP levels did not change significantly after cocaine treatment. * p < 0.05. Hippocampus: Saline n = 9, Cocaine n = 9, Morphine n = 8; Striatum: Saline n = 9, Cocaine n = 9, Morphine n = 9.

Prior exposure to cocaine or morphine did not result in altered levels of mBDNF or proBDNF (Fig. 6) or in altered levels of total GSK3β, pGSK3β, or pGSK3β/total GSK3β (Fig. 7) in either hippocampus or striatum (for all ANOVAs, p’s > 0.2). However, regional differences in BDNF were observed, with tissue levels of both proBDNF and mBDNF significantly higher in the hippocampus than in the striatum (p’s < 0.0001; Fig. 6).

**Fig. 6.**
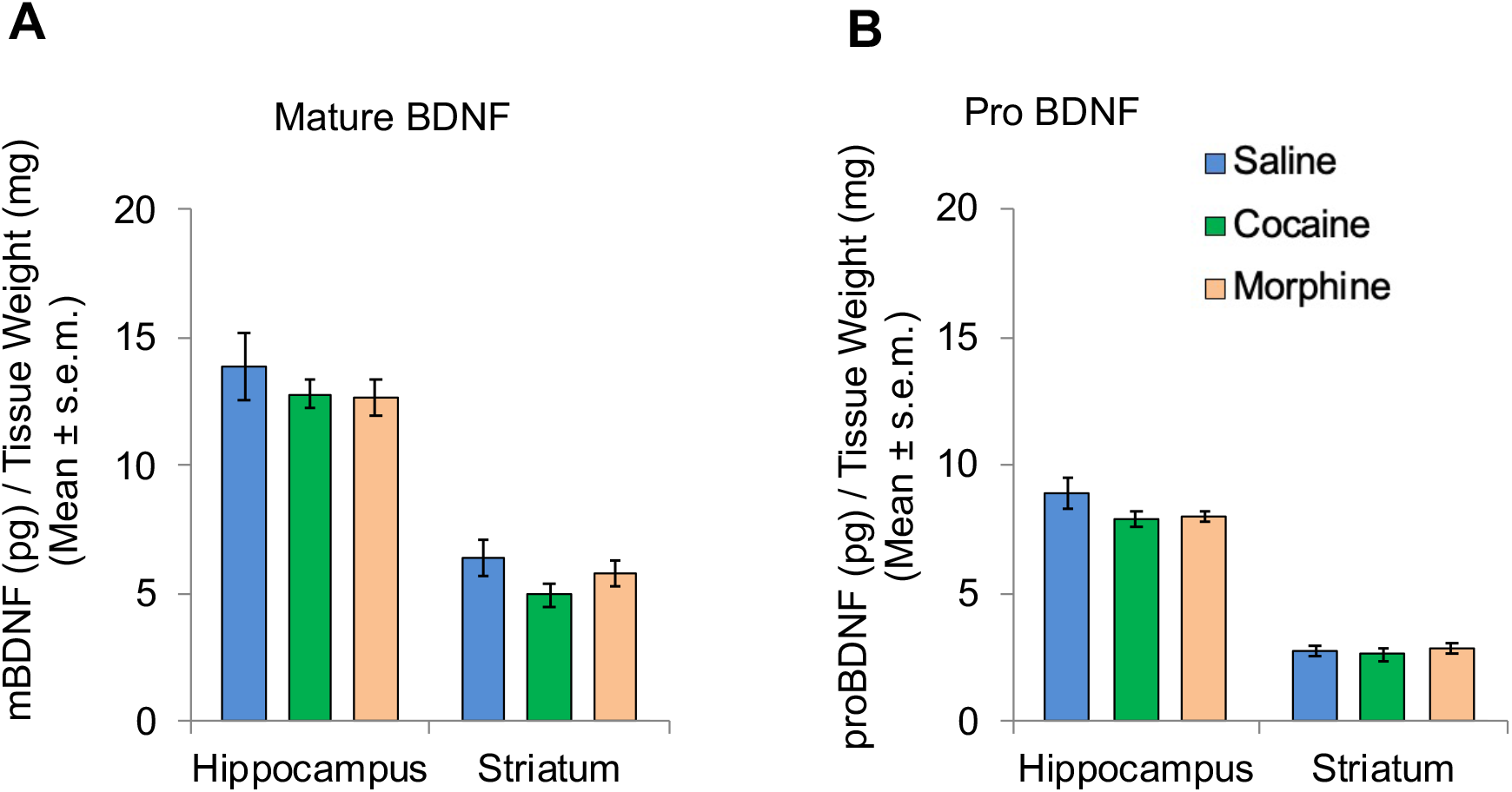
Levels of brain-derived neurotrophic factor (BDNF) (pg/mg wet weight) in the hippocampus and striatum one month after drug exposure as measured with ELISAs. Levels of (A) mature BDNF and (B) pro BDNF did not differ significantly by drug treatment in either brain area. Hippocampus: Saline n = 7, Cocaine n = 8, Morphine n = 6; Striatum: Saline n = 6, Cocaine n = 8, Morphine n = 7.

**Fig. 7.**
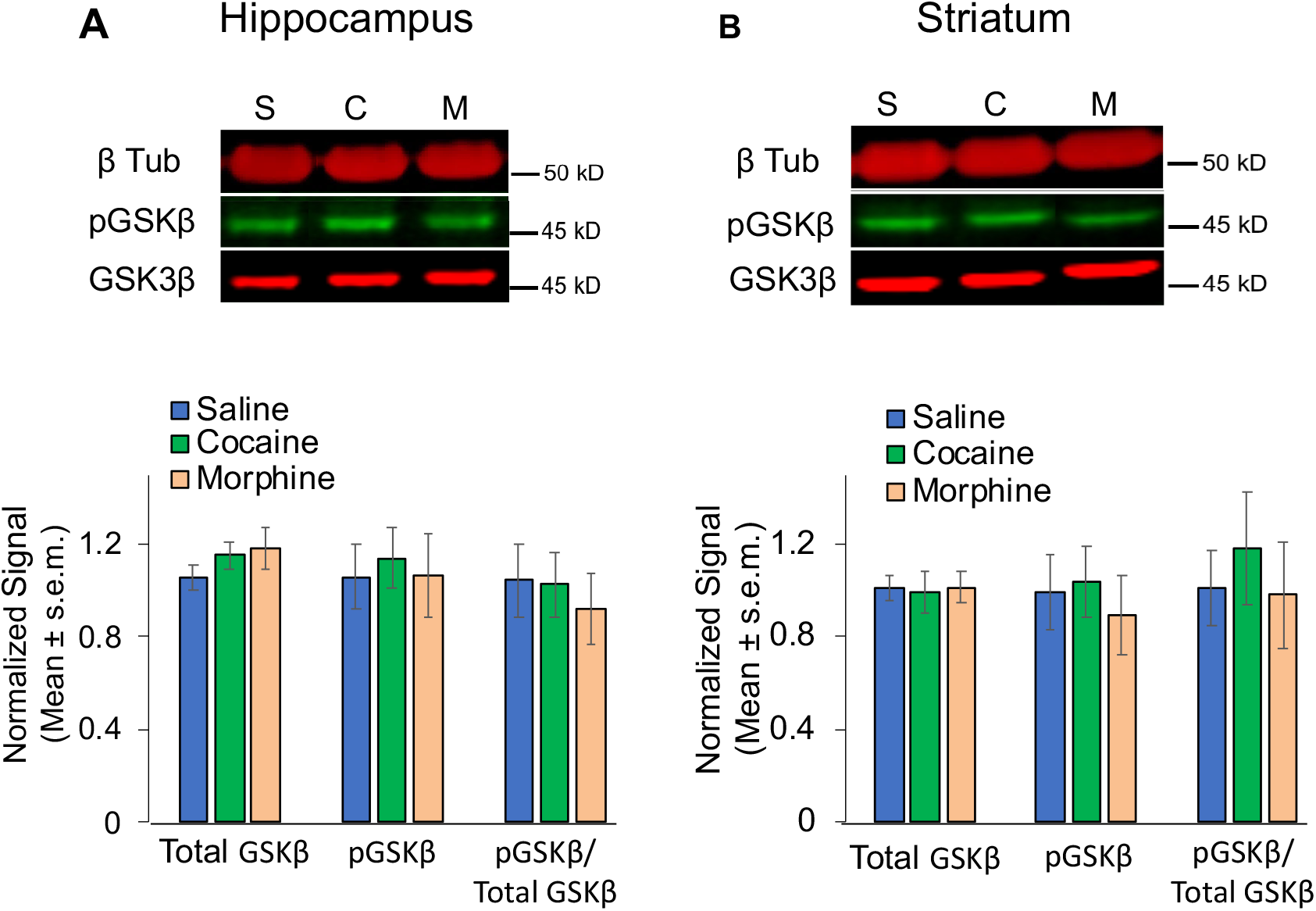
Levels of total and phosphorylated glycogen synthase kinase 3β (GSK3β) in the hippocampus and striatum one month after drug exposure as measured with western blots. (A) Mean levels of total GSK3β, phosphorylated (ser9) GSK3β, and the ratio of pGSK3β to total GSK3β in the (A) hippocampus and (B) striatum one month after drug treatments were not different across conditions. Representative bands are shown for each target. Hippocampus: Saline (S) n = 9, Cocaine (C) n = 9, Morphine (M) n = 9; Striatum: Saline (S) n = 9, Cocaine (C) n = 8, Morphine (M) n = 9.

## 4. Discussion

The findings reported here show that morphine treatment results in impaired learning on hippocampal-sensitive tasks and enhanced learning on striatum-sensitive tasks evident long after drug exposure. Because hippocampal and striatal contributions to cognition are often competitive, our findings of drug-induced changes in learning and decision-making could reflect impairments to processing within a hippocampus-based system, augmented processing within a striatum-based system, or both.

In single-solution tasks, 5-days of morphine exposure impaired place learning and enhanced response learning when tested after one month of abstinence. While place and response tasks differ according to the learning rule required to find food rewards, the reward itself, the apparatus, and motor demands are the same.

Importantly, performance on the tasks was at chance levels and comparable across drug groups early in training, suggesting that drug-effects on non-mnemonic aspects of the tasks, for example, on locomotion and motivation to explore, did not appreciably contribute to our findings. Thus, comparisons of learning on the two tasks provide a way to study the effects of prior drug use on learning and memory functions across memory systems.

Prior exposure to morphine also resulted in a robust shift away from the use of place (hippocampal) strategies and toward response (striatal) strategies in a dual-solution task that allowed rats to use either strategy. Importantly, prior drug exposure did not alter the speed of learning in the dual-solution task, but rather shifted the strategy or decision-making process used to solve the task. These findings are analogous to others using the dual-solution task to reveal changes in hippocampal (place) vs striatal (response) strategies during learning (Gardner et al., 2013, 2016, 2020a; Gold et al., 2013; Gold, 2016; Hawley et al., 2012; Korol and Wang, 2018; Korol et al., 2004; McElroy and Korol, 2005; Packard and Goodman, 2012, 2013; Packard, Goodman, and Ressler, 2018; Packard and McGaugh, 1996; Schwabe, 2013). The internal consistency of our findings across maze tasks provides strong evidence of a long-lasting drug-induced shift in memory system engagement during cognition that favors striatal control over behavior.

The present findings show that morphine induces long-lasting changes in learning and decision-making processes. Of interest, while cocaine generally produced behavioral results in the same direction as those seen with morphine, the results from cocaine-treated rats were not significantly different than those from saline controls for the learning tasks tested here. It is possible that cocaine and morphine differ in the long-term consequences of drug exposure on the balance of functions across memory systems. It is also possible that cocaine can support changes in behavior like those observed here with morphine, but that comparable effects of cocaine would emerge at doses or drug exposure regimens different than those included here.

Our findings are largely consistent with past reports using rats and mice showing morphine-induced impairments after periods of abstinence on learning and memory tasks that tap hippocampal functions (e.g., Dougherty et al., 1996; Harris and Aston-Jones, 2003; Shahroodi et al., 2020). Additionally, our findings show that a bout of morphine exposure elicits enhancement of learning and memory on striatum-sensitive tasks and reliance on striatum-based functions during decision-making long after the binge. A shift in learning from spatial to response strategies following substance abuse has been seen previously in short-term studies. For example, rats showed impaired spatial learning but enhanced response or cued learning after acute administration of alcohol in rats (Matthews et al., 1999; Sun et al., 2018). The shift in the selection of place or response strategies in rats is similarly seen in humans after drug abuse (e.g., Biernacki et al., 2016; Bohbot et al., 2013).

A lasting bias toward habit-based decision-making at the expense of goal-directed planning after drug exposure may consequently contribute to persistent maladaptive responses, e.g., drug-seeking without consideration of consequences, a hallmark of addiction (Lüsher et al., 2020; Milton and Everitt, 2012; Redish et al., 2008). The present findings suggest that changes to the balance of hippocampal vs. striatal functions following morphine treatment may underlie maladaptive choices observed in addiction. The shift from hippocampal to striatal processing is also seen in several other conditions including reproductive or estradiol status (Korol and Kolo, 2002; Korol et al., 2004; Korol and Pisani, 2015), aging (Barnes et al., 1980; Bohbot et al., 2012; Gardner et al., 2020b; Rodgers et al, 2012; Wiener et al., 2013), and stress (Packard and Goodman, 2013; Sadowsky et al., 2009; Schwabe, 2013). Of these factors, stress, in particular, has a prominent role in addiction and relapse, and stress may be an important mediator of the long-lasting changes across memory systems seen here. This idea is supported by examples of heightened stress and anxiety following withdrawal from drugs of abuse, including withdrawal from morphine, that can persist during prolonged abstinence (Alves et al., 2017; Welsch et al., 2020; also see Goodman and Packard, 2016; Schwabe et al., 2011).

Of the molecular markers assessed here, only GFAP showed changes consistent with the behaviorally measured shift from hippocampal to striatal processing after morphine exposure. In past reports, increases in GFAP protein or mRNA levels were associated with learning impairments on hippocampus-sensitive tasks in senescent rats (Sugaya et al., 1996) and in rats fed a high-fat diet (Bondan et al., 2019). Here, levels of GFAP in the hippocampus but not the striatum were increased one month after exposure to morphine and were thereby associated with depressed hippocampal functions. As stated above, stress too shifts the balance between hippocampal and striatal systems and may thereby contribute to the effects of drug abuse observed here. It is therefore of note that stress also modulates the expression of hippocampal GFAP (Jauregui-Huerta et al., 2010; Lambert et al., 2000).

The findings seen with GFAP measurements suggest that drug-induced alterations in hippocampal astrocytes may contribute to or may reflect shifts in hippocampal and striatal memory system functions. Consistent with our GFAP findings, several reports show acute effects of drugs of abuse on astrocytes in the hippocampus, including alterations in metabolic enzymes related to lactate production (Chen et al., 2007), in lactate transporters (Lindberg et al., 2019), and in glutamate transporters (Ozawa et al., 2001). Importantly, astrocytic regulation of these processes is also linked to learning and memory functions across systems (Alberini et al., 2018; Gold et al., 2013; Korol et al., 2019; Newman et al., 2017).

The results that mBDNF, proBDNF, and GSK3β levels or activation states remained unchanged one month after morphine or cocaine exposure contrast with those of prior reports showing alterations in BDNF and GSK3β levels in numerous brain regions, including the hippocampus and striatum, following drug exposure (e.g., Li and Wolf, 2015; Miller et al., 2014; Shahroodi et al., 2020). We have previously detected significant changes in these targets following training (Korol et al., 2013); thus the discrepant results likely reflect methodological differences in general experimental design across studies and not our ELISA or western blotting techniques.

In conclusion, the present report used a multiple memory systems approach to examine long-term consequences of morphine and cocaine exposures on learning and memory. Drug-induced shifts in cognitive style reflected in impairments in hippocampal learning, enhancements in striatal learning, and biases for striatum-based solutions during decision-making were most strikingly evident after prolonged abstinence from morphine. Prior morphine exposure also produced alterations in hippocampal astrocytes. Future work to evaluate a role for select functions of astrocytes in drug-related shifts across memory systems may identify novel targets to combat the persistent cognitive consequences of drug abuse.

## Acknowledgements

This work was supported by NIDA DA038798, NSF IOS 13-18490, NIA AG057947, and by P30 AG034464 through the Center on Aging and Policy Studies at Syracuse University.

